# Comparing DNA Extraction Protocols for Freshwater Prokaryotic Communities: Impacts on Yield and Microbial Profiling

**DOI:** 10.64898/2026.01.21.699709

**Authors:** Arthur Szylit, Ludwig Jardillier, Maria-Cristina Ciobanu, Léa Cabrol, Maialen Barret, Urania Christaki

## Abstract

DNA extraction from aquatic samples is a critical process that influences the quantity and purity of the DNA obtained. This can have profound effects on the accuracy of the community depiction. In this study, DNA extraction workflows of seven commercial kits (Qiagen: DNeasy PowerLyzer PowerSoil, DNeasy PowerSoil Pro, DNeasy PowerSoil, DNeasy PowerMax Soil; Macherey-Nagel: NucleoSpin Soil; Zymo: ZymoBIOMICS DNA; MP Biomedicals: FastDNA SPIN), along with several modifications of manufacturer’s protocols focusing on the lysis and elution steps, were tested, accounting for a total of 18 different protocols. For each protocol, DNA yield (quantity, replicability and quality), richness and compositional reproducibility based on 16S rRNA gene sequencing, as well as processing time and cost were assessed. The standard protocols recommended by the manufacturer showed comparable DNA yield results. Shared ASVs between all protocols accounted for >90% of the reads and were mostly abundant ASVs, indicating consistent detection of dominant taxa across all protocols. Adding supplementary lysis and elution steps to the manufacturer’s protocols yielded up to ∼4× more DNA. However, total read counts and ASV richness were lower as total DNA increased. Manufacturer’s protocols therefore showed higher values than their modified versions, although these effects were not significant on community composition. We conclude that the choice of a protocol is the balance between recovering sufficient DNA of good quality versus potential effects on downstream sequencing output (reads and ASVs).

## Introduction

No less than three centuries separate Leuwenhoek’s pioneering observations of microorganisms and the direct enumeration of aquatic bacteria [1]. It is well known that less than 1% of natural bacteria can be cultured and studied under standard laboratory conditions, a phenomenon called “the great plate count anomaly” [2–4]. Our knowledge of the composition and functions of microbial communities from a variety of habitats on Earth has expanded greatly over the past decades, driven primarily by progress in DNA sequencing technologies [5,6] to a point that environmental science has moved into the ‘age of the microbes’ [7]. Accurately deciphering community composition and function is essential for addressing key ecological questions, such as understanding how microbial communities are structured in space and time, how this structure affects their functional roles, and ultimately their contribution to biogeochemical cycles. The reduction in both cost and processing time of high-throughput sequencing platforms facilitated their widespread adoption [8,9]. Yet the accuracy of sequencing-based analyses depends strongly on upstream steps, particularly the isolation of high-quality DNA from environmental samples and, for metabarcoding approaches, the choice of primer sets [10]. DNA extraction from aquatic samples is a critical but often delicate process influencing DNA quantity and purity that can have profound effects on the accuracy of the community composition assessment [11]. Despite initiatives for standardization in the framework of international projects such as the Earth Microbiome Project (https://earthmicrobiome.org/protocols-and-standards/), Ocean Sampling Day Initiative [12], NEON [13] and Biogeoscapes (https://biogeoscapes.org/si/), no consensus method has been adopted so far. Differences in sample origin, targeted organisms, research goals, and available resources often lead laboratories to adopt protocols best suited to their constraints and objectives. Historically, DNA extraction relied on phenol-chloroform protocols, combining enzymatic or chemical lysis with ethanol precipitation [14]. Although effective, these methods used hazardous chemicals, were time-consuming, and difficult to standardise. The growing interest in microbial research has led to the development of numerous commercial kits. While the exact composition of these kits is generally not disclosed for commercial reasons, they typically follow similar steps: cell lysis, DNA binding, washing, and elution. Despite these common steps, several studies have reported significant differences between manufactured kits in terms of DNA yield and the downstream analysis of prokaryotic communities across diverse sample types including, for example, soils [15], permafrost [16,17], bioreactors [18], marine sediments [19], plant tissues [20], human microbiomes [21,22], ancient DNA [23], and freshwaters [24].

In aquatic ecosystems, analysing prokaryotic communities typically involves a filtration step to concentrate biomass onto a membrane. While this filtration step, through filter porosity and subsequent treatments, significantly influences community differentiation [25], high levels of suspended particulate matter —independently of prokaryote abundance—can quickly clog filters, thereby reducing microbial biomass recovery and impairing sample quality. Such conditions are common in estuarine and coastal marine environments, and also occur in most freshwater ecosystems. Limited size and depth make freshwater ecosystems particularly sensitive to allochthonous inputs such as erosion runoff, wind and rainfall, which can load particles or resuspend previously settled particles. In this context, the DNA extraction step of water samples remains critical, as it aims to recover as much DNA as possible from a potentially low biomass. The objective of the present study was to provide practical recommendations for selecting and optimizing DNA extraction workflows for prokaryotic community studies in freshwaters that, by extension, could also apply to microbial eukaryotes and marine environments. Seven commercial DNA extraction kits were tested. In addition to manufacturers’ protocols, we benchmarked targeted lysis and elution modifications, leading to 18 protocols that we compared. DNA recovery efficiency was evaluated in terms of DNA yield, richness and compositional reproducibility using 16S rRNA gene sequencing, as well as processing time and cost.

## Material and Methods

### Sample collection

An artificial lake located near Paris, France (48.715506° N, 2.157887° E) was sampled in 2023. This 330 m² waterbody was formed in 2017. Water samples were collected from the shoreline using a clean plastic beaker attached to a 3 m rod and transferred into a sterile 10 L container. The collected water was prefiltered sequentially through 100-µm and 30-µm polycarbonate membranes (Milipore Isopore) to remove large particles. The prefiltered water was filtered onto a 0.2-µm pore-size and 47 mm diameter polycarbonate membrane (Whatman Nucleopore). The same volume (200 mL) was filtered for all samples. A total of 57 filters were collected across two independent sampling events, each carried out on a single day: Experiment K (Kit comparison, 45 filters) in January and Experiment L (Lysis optimization, 12 filters) in September. As the two experiments corresponded to distinct events, results were compared only within each experiment and never between them. All membranes were immediately placed in sterile Petri dishes and stored at –80 °C until further processing, between 3 days and 1 month after collection.

### DNA extraction protocols

We chose to test some of the most commonly used manufacturer DNA extraction kits. The kits and associated protocols that were tested are summarised in Table 1. Originally designed for soil matrices, these kits include more stringent purification steps than standard water protocols, and are therefore more appropriate for inhibitor-rich samples such as those usually collected in freshwaters. For all tests, membranes were cut with scissors sterilised by ethanol and flaming under aseptic conditions prior to DNA extraction to enhance the efficiency of bead-based mechanical lysis. All subsequent extraction procedures were performed on ice in triplicate to assess technical reproducibility and enable statistical comparisons between protocols.

**Table 1:**
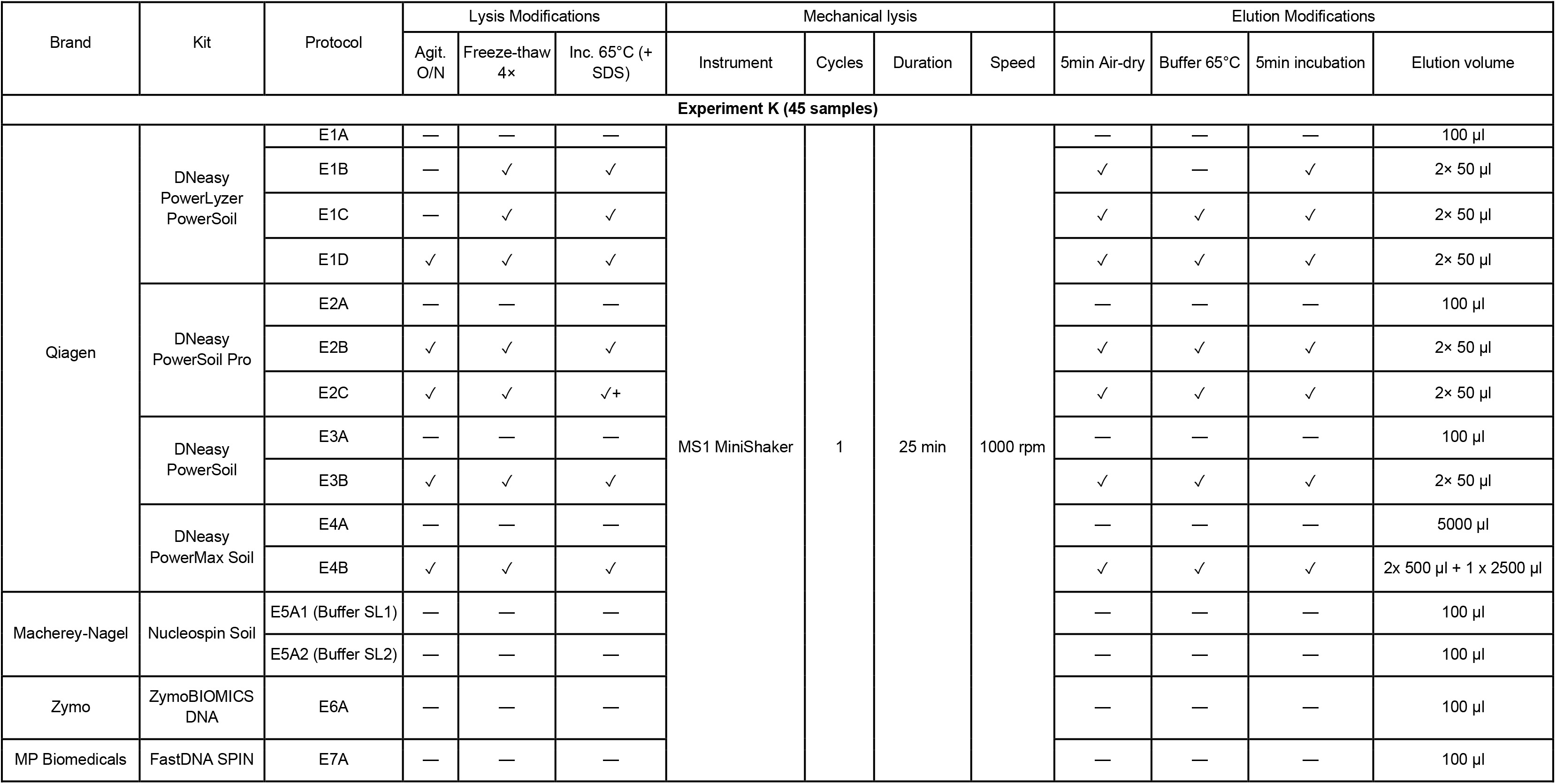
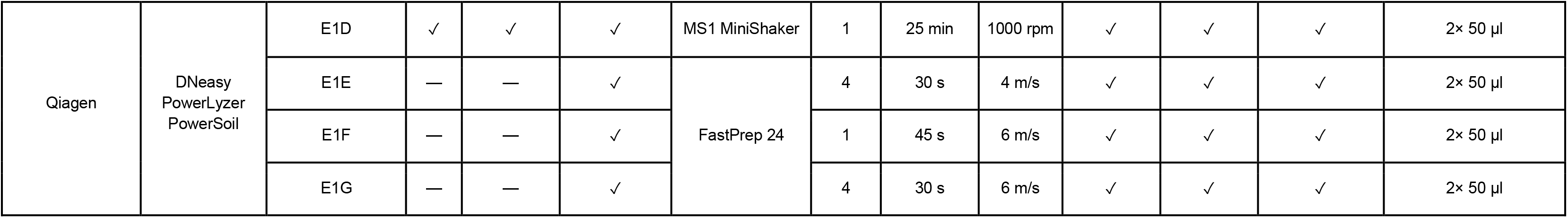
Summary of DNA extraction kits and protocols tested during the two experiments. Protocols E1–E7A correspond to the manufacturers’ protocols, while other codes indicate modified versions. Checkmarks (✓) indicate applied modifications; dashes (—) indicate omissions. The “+” denotes SDS addition during 65 °C incubation. In the “Elution volume” column, the notation “n× X µl” indicates n sequential elution steps of X µl each.

Experiment K (Kit comparison) compared 15 extraction protocols derived from seven commercial kits (E1 to E7) produced by four different manufacturers (Table 1). For each kit, the manufacturer protocol was followed (indicated by suffix “A”, Table 1). For the NucleoSpin Soil kit (Macherey-Nagel), E5A1 and E5A2 correspond to the manufacturer’s protocol using lysis Buffers SL1 and SL2, respectively (Table 1). Modified protocols for Qiagen kits (E1D, E2B, E3B, E4B) incorporated the following variations (Table 1): membranes were first incubated overnight at 5°C with agitation using a HulaMixer (Thermo Scientific™ 15920D) in PowerBead Solution (or CD1 for DNeasy PowerSoil Pro). For the PowerLyser kit, two simplified protocols (E1C and E1B) omitted this overnight incubation step. Following this, samples underwent four freeze-thaw cycles between liquid nitrogen (until completely frozen) and a 65°C water bath (until fully thawed). For PowerSoil, PowerMax, and PowerLyser PowerSoil kits, Solution C1 was then added and samples were incubated at 65°C for 10 min prior to mechanical lysis. For the DNeasy PowerSoil Pro (which does not use Solution C1), protocols were tested with (E2C) and without (E2B) the addition of 80 µL of 10% SDS (final concentration 1%) before the 65°C incubation. Mechanical lysis was performed by vortexing for 25 min at 1000 rpm (Table 1). Elution modifications included extended column drying by placing tubes on ice with loose caps for 5 min following centrifugation. Elution was performed in two sequential steps using preheated elution buffer at 65°C (room temperature for E1B): 50 µL was added, incubated for 5 min, and centrifuged; this step was repeated with an additional 50 µL. For the PowerMax kit, the manufacturer’s protocol (E4A) uses a single 5 mL elution, whereas the modified protocol (E4B) employed three sequential elution steps (500 µL, 500 µL, and 2500 µL; Table 1)

Furthermore, experiment L (Lysis optimization) compared four extraction protocols using the PowerLyser kit, all starting from the E1D protocol. E1D was retained as a control, while three modified versions (E1E, E1F, E1G) replaced the overnight incubation, freeze-thaw cycles, and vortex-based mechanical lysis with FastPrep-24 bead-beating (MP Biomedicals). FastPrep conditions varied in cycle number, duration, and speed as specified in Table 1. In protocols with multiple cycles (E1E and E1G), a 30-second pause was included between cycles to reduce heat buildup. All other steps (Solution C1 incubation, elution modifications) remained identical to E1D.

DNA quality assessment was measured for all extractions from both experiments, where DNA fragment size was assessed on a 1% agarose gel, and absorbance ratios (A260/280 nm and A260/230 nm) were measured using a NanoDrop 2000 spectrophotometer (Thermo Scientific). The A260/280 ratio reflects protein contamination, with values around 1.8 considered indicative of pure DNA, whereas the A260/230 ratio reflects contamination by organic compounds and salts (e.g. carbohydrates, phenolic compounds, guanidine), with optimal values around 2.0–2.2. DNA quantity was determined using a Qubit 3 Fluorometer with the Qubit dsDNA BR Assay Kit (Invitrogen)

### 16S rDNA amplification and sequencing

Prokaryotic community composition of all samples (57 samples) was determined by sequencing the 16S rRNA gene in prokaryotes (bacteria and archaea), using primers U515F and Pk926R (411 bp) targeting the V4–V5 region [26], following the Earth Microbiome Project protocol (https://earthmicrobiome.org/protocols-and-standards/16s/). PCR amplification was carried out using Platinum Taq DNA Polymerase (Invitrogen, Thermo Fisher Scientific, Waltham, MA, USA), following the manufacturer’s recommended reaction mix (Table S1), supplemented with DTT from the PCR Decontamination Kit (Enzo Life Sciences) as per manufacturer’s instructions. A fixed volume of 1 µl of DNA extract was added to each reaction, regardless of its concentration. Thermocycling consisted of an initial denaturation at 94 °C for 2 min, followed by 35 cycles of 94 °C for 15 s, 55 °C for 30 s and 72 °C for 1 min 30 s, a final extension at 72 °C for 5 min, and a hold at 10 °C. Amplicon length was verified by electrophoresis on a 1% agarose gel. For each sample, products from 3 to 5 PCR replicates were combined and purified using the QIAquick PCR Purification Kit (Qiagen, Germany). Amplicon concentrations were normalised to 20 ng.μL^-1^ prior to sequencing. All libraries were sequenced in a single MiSeq 2 × 250 run (GENEWIZ, Germany) to avoid inter-run bias.

### Bioinformatic pipeline

A total of 2 806 613 raw paired-end sequences were processed using the DADA2 pipeline [27] implemented in R (version 4.3.3). Reads were filtered and trimmed using the *filterAndTrim()* function with the following parameters: forward and reverse reads were truncated at 240 and 200 bases, respectively, based on quality profiles generated with QIIME 2 version 2023.9 [28], which showed a median Phred score of 38 at both positions. The first 19 and 20 bases were removed to eliminate primers, ambiguous reads were discarded (maxN = 0), and low-quality tails were trimmed at a quality threshold of 2 (truncQ = 2). Error rates were learned with the *learnErrors()* function, and amplicon sequence variants (ASVs) were inferred using *dada()*. Read pairs were merged using the *mergePairs()* function with a minimum overlap of 15 bp and a maximum of 1 mismatch. Only merged sequences between 364 and 382 bp in length were retained. Chimeric sequences were removed using the *removeBimeraDenovo()* function with the consensus method. ASV taxonomic assignment was performed with the *assignTaxonomy()* function, using the naive Bayesian classifier trained on the SILVA SSU reference database version 138.1 [29], with reverse complement detection enabled (tryRC = TRUE). Before analysis, unclassified ASVs at the phylum level, or affiliated with Mitochondria or Chloroplast were removed. Singleton ASVs and those representing less than 0.005% of total reads across all samples were excluded [30], reducing the dataset from 3 709 to 371 ASVs (about 10% retained) while preserving 98.4% of the reads (from 1 856 016 to 1 826 234 reads). The sequence dataset was then rarefied to the minimum sequencing depth observed across samples (19 485 reads), resulting in a final dataset of 57 samples, 1 110 645 sequences, and 371 ASVs. All raw sequences have been deposited in the European Nucleotide Archive under project accession number *PRJEB94317*.

### Data Analysis

All statistical analyses were performed using R version 4.3.3 [31]. To compare DNA yield, DNA quality and alpha diversity indices between extraction conditions, a non-parametric Kruskal-Wallis test was used, The prokaryotic dataset was processed using the phyloseq package version 1.46.0 [33]. Data manipulation was performed using the dplyr package version 1.1.4 [34] and tidyr version 1.3.1 [35]. Graphs were created using the ggplot2 package version 3.5.1 [36]. Alpha diversity indices (Richness, and Evenness) were calculated from the rarefied prokaryote dataset using the *estimate_richness* function within the phyloseq package for each sample. The core community was defined as the set of ASVs common to all samples (100% prevalence), with a relative abundance of at least 0.001%. Rare ASVs were defined as those with a relative abundance below 0.01%. A phylogenetic tree was constructed to compute Weighted UniFrac distances. Reference sequences were aligned using DECIPHER [37], and a Neighbor-Joining tree was built from maximum likelihood distances with phangorn [38]. The tree was then optimised under a GTR+I+Γ model using nearest-neighbor interchange (NNI) rearrangements. Weighted Unifrac distance [39] was used to study beta diversity in the rarefied prokaryotic community. A Principal Coordinate Analysis (PCoA) was performed to visualise the structure of the prokaryotic communities between samples. Differences in prokaryotic community composition between extraction methods were assessed using a permutational multivariate analysis of variance (PERMANOVA) [40] with the *adonis2()* function from the vegan package (version 2.6.4) [41], after verifying the homogeneity of multivariate dispersions using the *betadisper()* function (vegan package). Differential abundance analysis was performed at the genus level using ANCOM-BC [42] to identify genera that significantly differed in abundance according to the extraction protocol (independently for each sampling campaign). P-values were adjusted for multiple testing using false discovery rate (FDR) correction. Only genera with an adjusted p-value below 0.05 and an absolute log₂ fold change greater than 1 were considered significantly differentially abundant.

## Results

### DNA yield (quantity and replicability)

In Experiment K, DNA yield ranged from 58.6 ± 24 ng for standard Nucleospin (E5A2) to 1745.7 ± 119.2 ng for modified DNeasy Powerlyser (E1D; Fig 1A). Although the NucleoSpin protocols (E5A1 and E5A2) yielded the lowest average amounts of DNA, this reduction was not statistically significant compared to the other unmodified manufacturer protocols (Kruskal-Wallis, p > 0.05).

**Figure 1:**
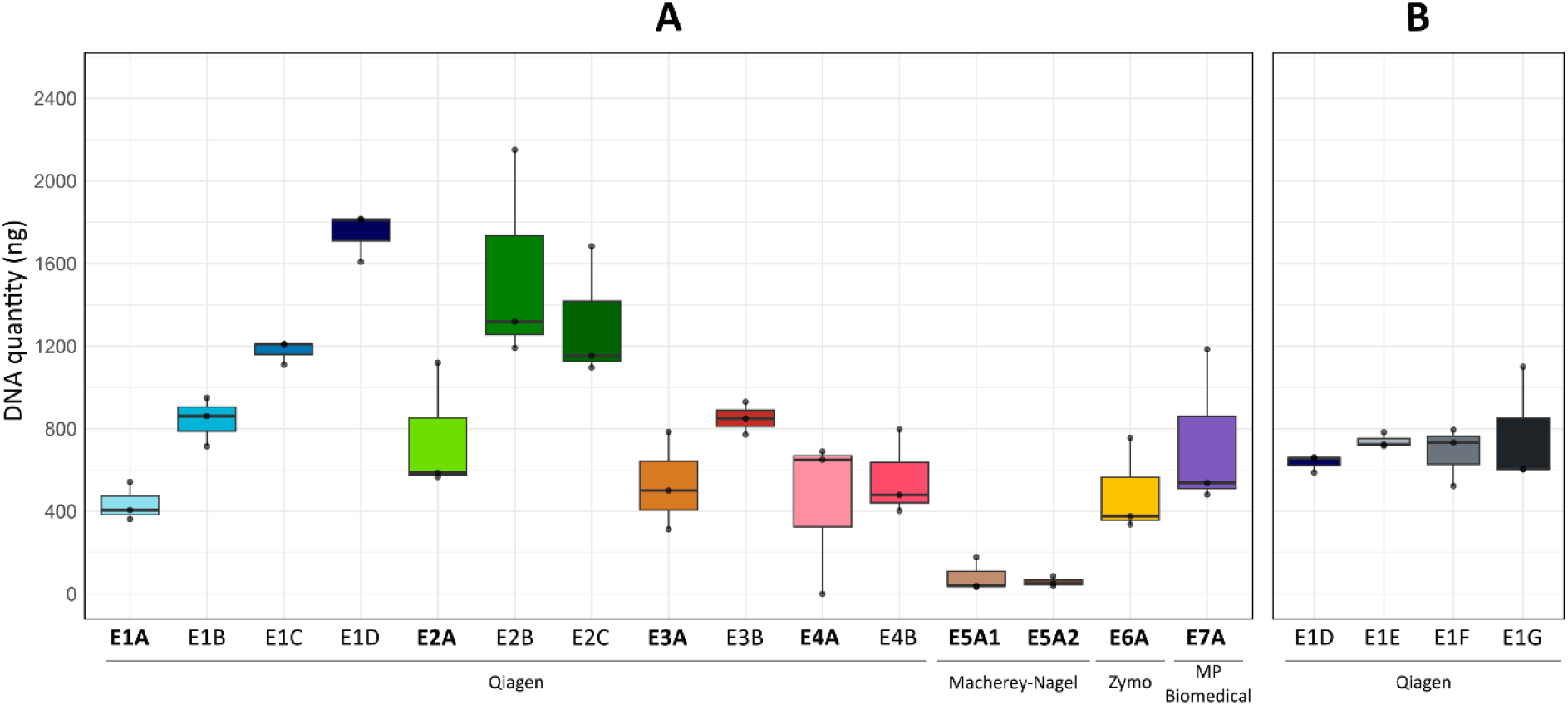
DNA yield across extraction protocols for Experiment K (A) and Experiment L (B).

For the four Qiagen kits, only the DNeasy PowerLyzer showed significant differences among protocols (Kruskal-Wallis, p = 0.02).. For the DNeasy PowerSoil Pro, the SDS addition in E2C did not increase yield relative to E2B. Overall, the protocol E1D combined the highest DNA yield (approximately 4-fold higher than the manufacturer’s protocol) with a low coefficient of variation (RSD=6.8%), compared with other modified protocols E2B and E2C (RSD=33.5% and 24.5%), indicating more consistent performance across replicates (Fig 1A).

Consequently, the DNeasy PowerLyzer E1D protocol (Table 1) was tested with stronger mechanical lysis in Experiment L. DNA yields did not then differ significantly among protocols E1D, E1E, E1F, and E1G (Kruskall-Wallis, p = 0.58; Fig 1B). E1D showed a coefficient of variation (CV) of 6.5%, while replicate variability increased with lysis intensity across the FastPrep conditions, with CVs of 4.9%, 20.8%, and 37.3% for E1E, E1F, and E1G, respectively.

### DNA quality

In Experiment K, absorbance ratios varied among protocols (Fig 2A). For A260nm/230nm, only PowerLyzer protocols (E1) showed significant differences between manufacturer and modified (Kruskall-Wallis, p = 0.03), with ratios of 0.9 ± 0.2 for the manufacturer protocol (E1A) and 1.6 ± 0.1 for the modified protocol (E1D) (Fig 2A). For A260nm/280nm, only PowerSoil Pro based protocols (E2) showed a difference (Kruskall-Wallis, p = 0.05), with ratios of 2.1 ± 0.1 for E2A and 1.9 ± 0.05 for E2C (Fig 2A).

**Figure 2:**
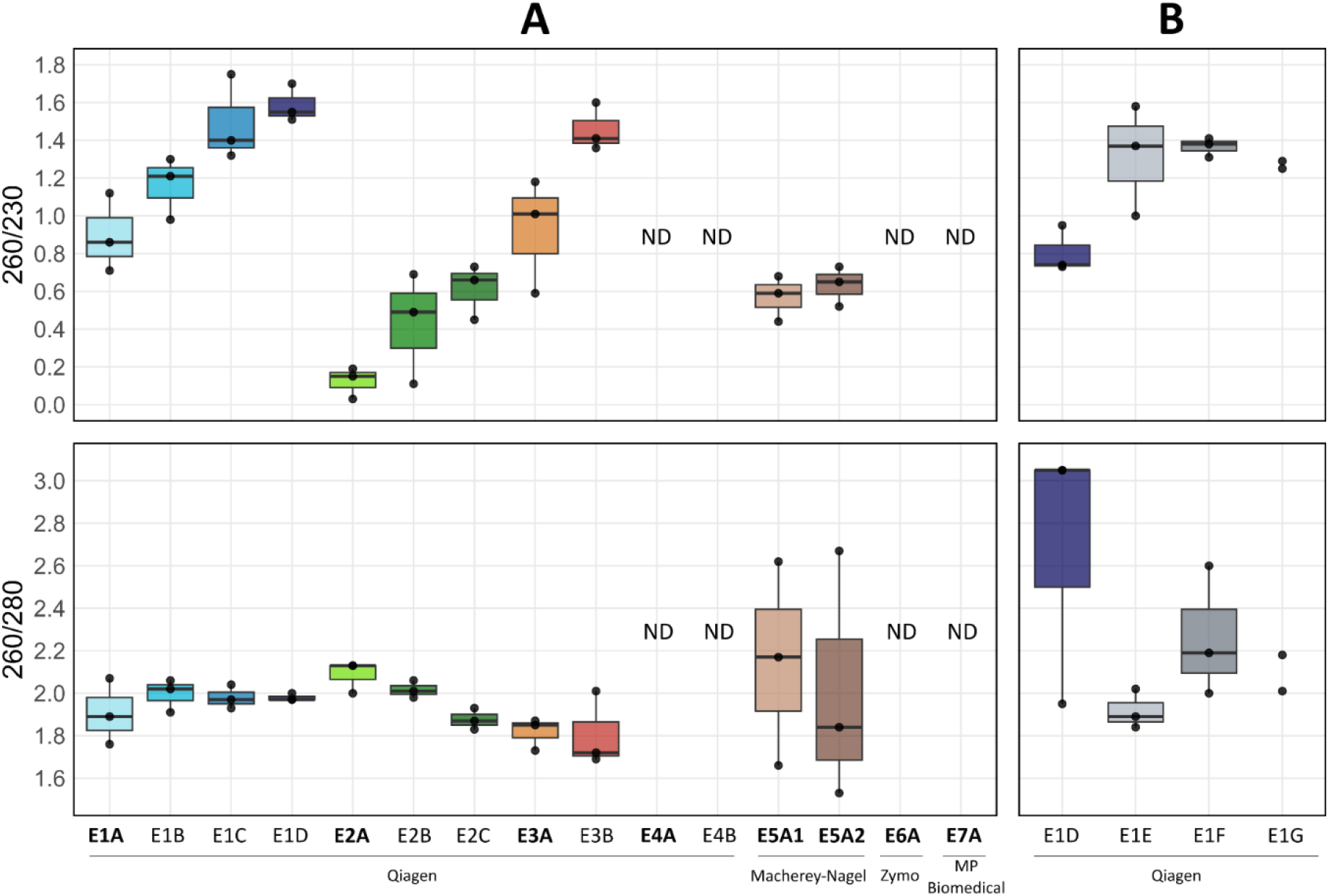
Absorbance ratios A260/230 and A260/280 across extraction protocols for Experiment K (A) and Experiment L (B). Protocols in bold correspond to manufacturers’ protocols. ND indicates aberrant or out-of-range NanoDrop ratios, resulting from low DNA concentration and/or co-extracted contaminants, despite sufficient total DNA measured by Qubit.

In Experiment L, A260nm/230nm ratios did not differ significantly among protocols (Kruskal-Wallis, p > 0.05), although FastPrep protocols (E1E/F/G) showed higher average ratios (1.32 ± 0.17) compared to E1D (0.81 ± 0.12; Fig 2B).

A260nm/280nm ratios did not differ significantly (Kruskall-Wallis, p = 0.28). Gel electrophoresis revealed differences in DNA fragment size between lysis methods (Fig S1): protocols without FastPrep-based lysis (E1D) produced DNA fragments of approximately 50 kb, whereas FastPrep protocols (E1E, E1F, E1G) yielded fragments ranging from 10 to 15 kb.

### Alpha diversity

In Experiment K, ASV richness ranged from 173 ± 7 (E2B) to 192 ± 7 (E1A) (Fig 3A). While the DNeasy PowerLyzer standard protocol (E1A) showed higher average richness compared to its modified versions (E1B, E1C, E1D), this difference was not statistically significant (Kruskal-Wallis, p = 0.08). Read counts mirrored this pattern, decreasing from 32 980 ± 3 189 for E1A to 28 683 ± 1 533 for E1D (Fig S2). Evenness did not differ significantly among manufacturer protocols (Kruskal-Wallis, p = 0.054), although the Macherey-Nagel kits (E5A1 and E5A2) numerically displayed the highest values (0.830 ± 0.004). For all Qiagen kits, manufacturer and modified protocols did not differ significantly for this index (Fig 3A). In Experiment L, ASV richness did not differ among E1D, E1E, E1F, and E1G (Fig 3B). Within E1F, two of the three replicates had lower ASV abundance (118 and 136 ASVs), yet the mean of the replicate values did not differ from the others (Kruskall-Wallis, p=0.08). Evenness differed significantly among the four protocols (Kruskal-Wallis, p = 0.02, Fig 3B), with E1G showing the lowest values (0.54 ± 0.02).

**Figure 3:**
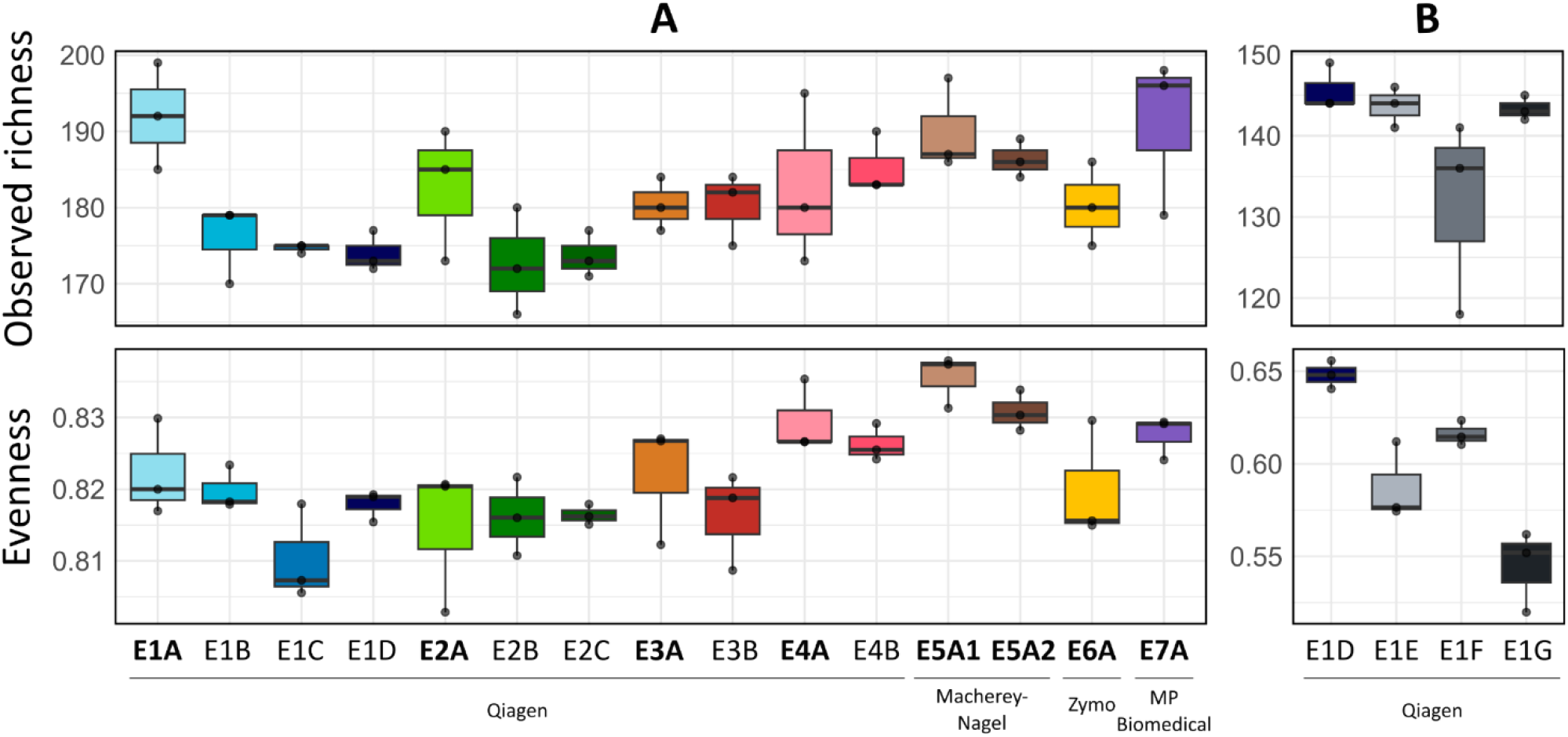
Alpha diversity indices (ASV richness and Evenness) of prokaryotic communities across extraction protocols for Experiment K (A) and Experiment L (B). Protocols in bold correspond to the manufacturers’ protocols.

Overall, taking in account all protocols and replicates, DNA yield was negatively correlated with ASV richness and reads number in experiment K (Pearson correlation, p< 0.001, Fig 4A-B).

**Figure 4:**
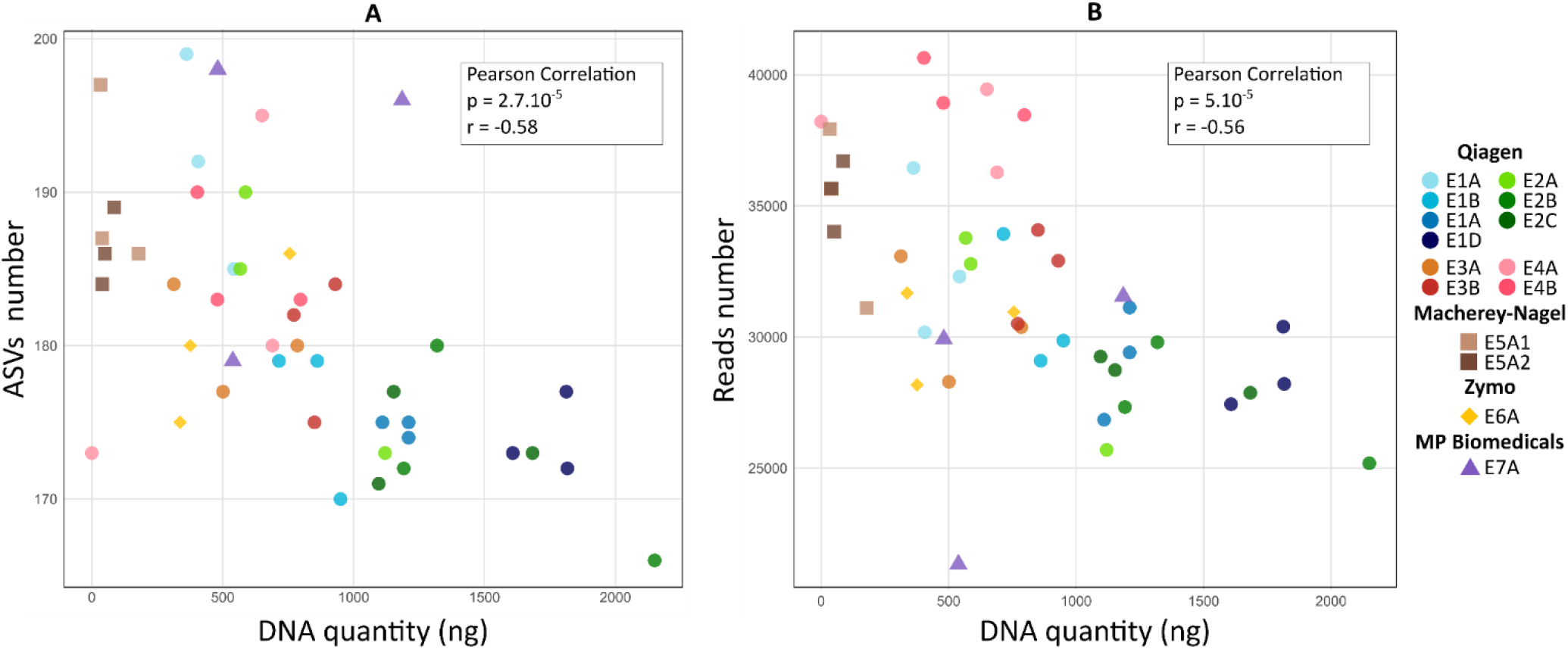
Relationship between DNA yield and sequencing output across extraction protocols in Experiment K. Correlation between DNA quantity and ASV richness (A) and between DNA quantity and the number of sequencing reads (B). Colors and shapes indicate the extraction protocol. Pearson correlation coefficients and p-values are shown in each panel.

### Community structure

In Experiment K, 109 ASVs (representing 42% of the total 261 ASVs) formed the core community (Fig 5A). On average, these ASVs accounted for 93.7 ± 1% of the reads per tested protocol. Similar results were obtained in Experiment L, where the core community comprised 96 ASVs representing 55% of the total 176 ASVs and accounted for 96.7 ± 0.8% of reads across protocols (Fig 5B). In Experiment K, most abundant core genera *Limnohabitans*, “hgcI clade” and “CL500-29” accounted for about 21.0% of the total reads. In Experiment L, the order *Synechococcales*, exclusively composed of the genus *Cyanobium*, was dominant in the core community across all protocols (38.1 ± 8.2%). Its relative abundance varied among protocols: from 28.5% (E1D) to 35.2% (E1F), 40.9% (E1E), and 47.7% (E1G) (Fig S3). In contrast to the core community, the number of rare ASVs varied among protocols (Fig S4A). In Experiment K, rare ASVs ranged from 14 (E6) to 25 (E4B). Manufacturer protocols generally detected more rare ASVs than their modified counterparts, except for PowerSoil (E3), which detected 18 in both versions. In Experiment L (Fig S4B), a single, intense FastPrep cycle (E1F) detected only 3 rare ASVs, compared to 7–8 in protocols using multiple cycles (E1E, E1G) or vortex-based lysis (E1D).

**Figure 5:**
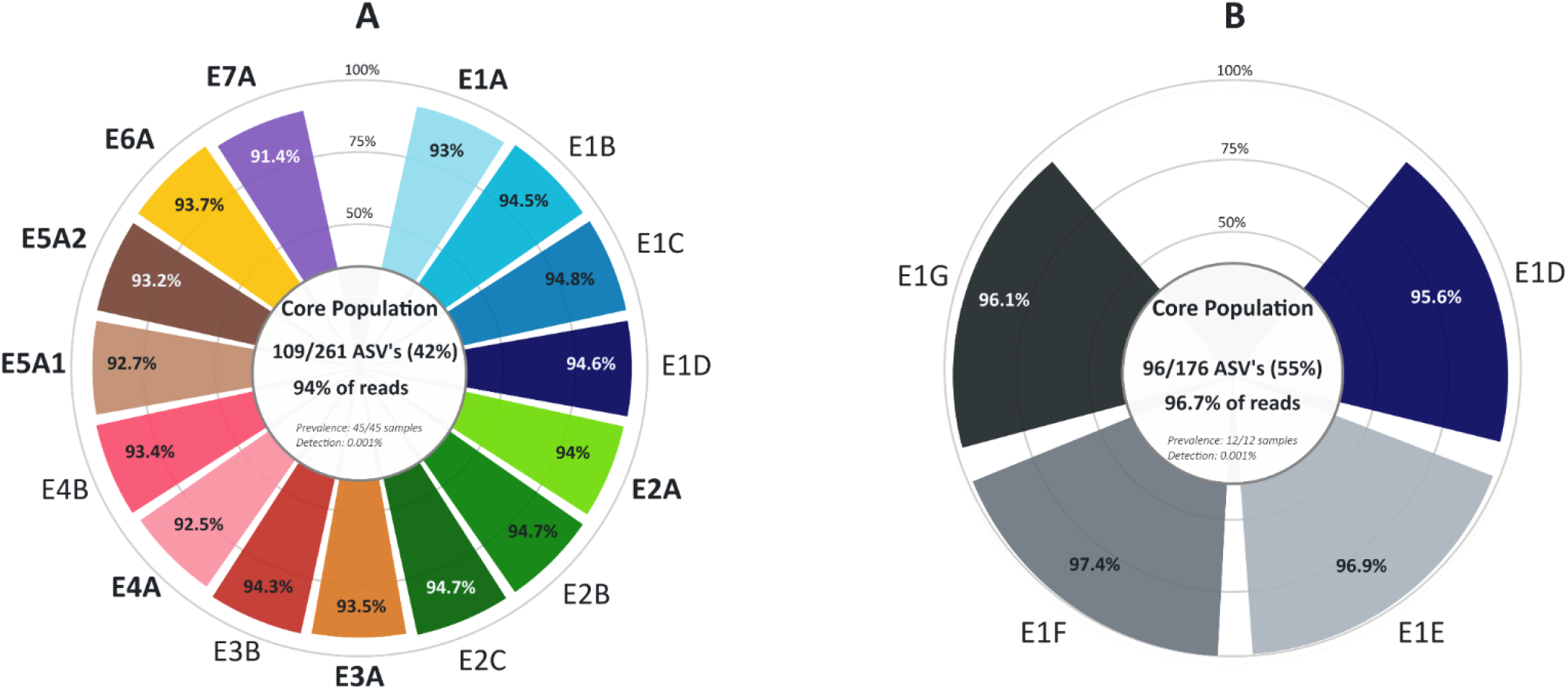
Core prokaryotic community across extraction protocols for Experiment K (A) and Experiment L (B). The core was defined using a prevalence threshold of 100% and a detection threshold of 0.001%. For each donut chart, the center shows the number and percentage of core ASVs and their corresponding reads, while the outer ring displays the proportion of reads assigned to core ASVs for each extraction protocol.

PCoA based on all ASVs showed significant differences among protocols (PERMANOVA, p = 0.001 for both experiments) and among manufacturers (PERMANOVA, p = 0.003 for Experiment K; Fig 6A-B). However, pairwise comparisons did not identify significant differences between specific manufacturers and protocols, and no clear phylogenetic pattern emerged. In Experiment K, differential abundance analysis revealed that several genera were enriched in manufacturer’s protocols compared to their modified counterparts (Fig S5A). Specifically, *Clostridium* was most enriched in E1A (vs E1D), *Fictibacillus* in E2A (vs E2B) and E3A (vs E3B), and *Tumebacillus* in E4A (vs E4B). In Experiment L, differential abundance analysis showed that genera from the Proteobacteria phylum were enriched in E1E and E1G relative to E1D (Fig S5B). While 13 genera differed in abundance between E1D/E1E and E1D/E1G, only one genus differed significantly between E1E and E1G (Fig S5B).

**Figure 6:**
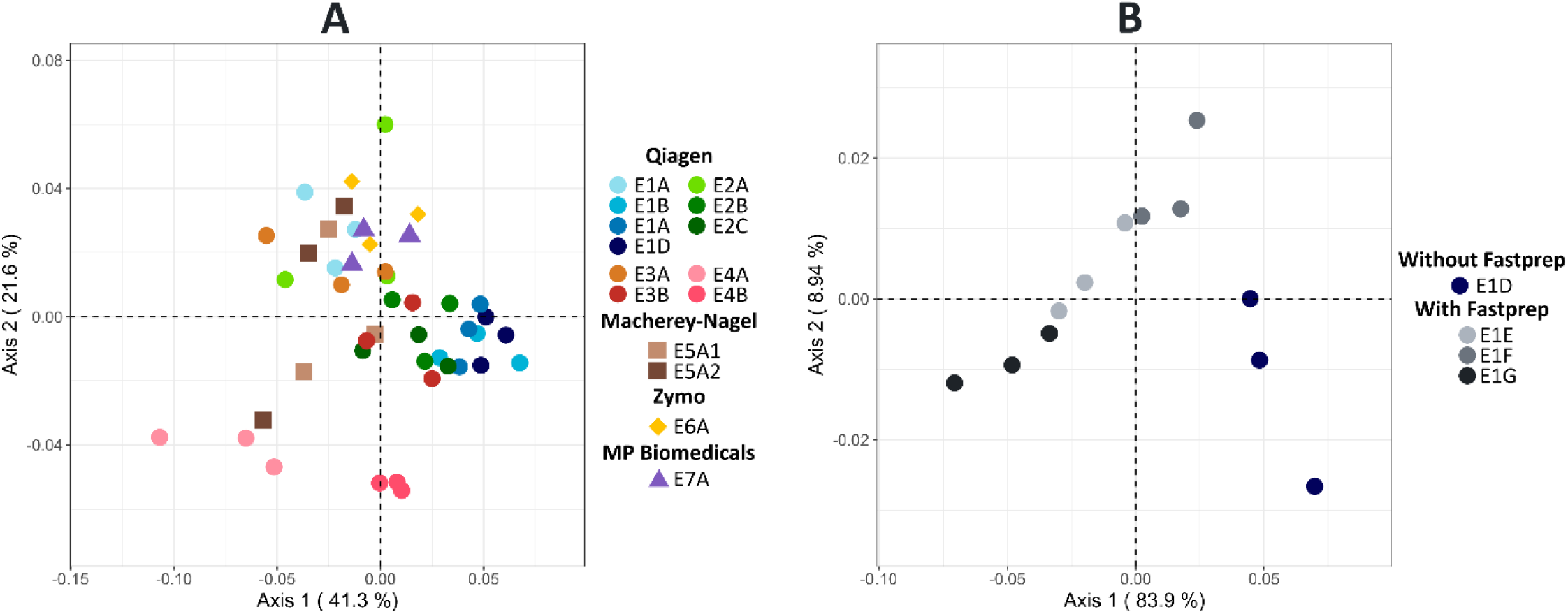
Principal Coordinate Analysis (PCoA) of prokaryotic communities for Experiment K (A) and Experiment L (B) sampling based on Weighted UniFrac distances. Each point represents a sample, colored and shaped according to the DNA extraction protocol used.

## Discussion

On a global scale, aquatic ecosystems contribute greatly to the functioning of the biosphere through their role in the regulation of climate and carbon flows, for example [43,44]. Elucidating the processes driving these ecosystems, particularly biogeochemical fluxes, is essential to appreciate their temporal dynamics and their responses to disturbances. Microbial communities drive all biogeochemical cycles and therefore are key players in the functioning of aquatic ecosystems. The development of molecular approaches and sequencing technologies has rapidly expanded our knowledge of the tree of life, mostly for microbial diversity. This advent has revealed the tremendous microbial phylogenetic diversity and its associated metabolism diversity. It is therefore of primary importance to be able to depict accurately how microbial community composition changes with their environment and, in consequence, to link any change in composition with changes in functions and ultimately on biogeochemical cycles. A large fraction of the microbial diversity consists of low-abundance taxa. Although traditional paradigms attributed the main roles in biogeochemical cycles to abundant taxa, numerous studies have emphasised the significant importance of rare taxa [45,46]. Accurately deciphering spatio-temporal changes in these taxa and their effects on biogeochemical cycles is essential. This requires careful evaluation of all analytical steps. While the effects of primer sets and bioinformatic pipelines have been extensively assessed [47,48], earlier steps, particularly DNA extraction, have received less attention.

Compared to previous studies, the present investigation assessed the differences in the DNA yield and quality as well as in the community depiction between a larger number of manufactured kits and protocol modifications (Table 2). As in previous works realised on freshwater ecosystems, most of the compared protocols involved Qiagen kits (Table 2). Qiagen kits compared here are routinely used for water, while Zymo and MP Biomedicals are more occasionally used [49,50]. The Qiagen DNeasy PowerSoil kit (E3, Table 1) has often been cited for high-quality DNA from water [24] and soil [16] and was later discontinued and replaced by the DNeasy PowerSoil Pro kit (E2). In the present study increasing DNA yield had a negative effect on community richness (Fig 4, Table 3). Although these effects were moderate, they demonstrate—as in previous studies (Table 2)—that protocol selection requires careful consideration of trade-offs based on study-specific objectives.

**Table 2:** Comparative studies of DNA extraction protocols from aquatic samples (2005-2023). The MoBio PowerSoil kit is the same as the Qiagen DNeasy PowerSoil kit.

| Study | Target organism | Matrix | Location | Methods compared | Manufacturers tested | Modification comparison | DNA Measurement | Target gene | Sequencing Approach | Main finding |
| --- | --- | --- | --- | --- | --- | --- | --- | --- | --- | --- |
| [75] | Bacteria Pathogens | Wastewater | Canada | commercial kit (1)<br>phenol-chloroform variants (6)<br>Other chemical methods (3) | MoBio | none | Yield Quality | 16S rRNA | targeted | Few of ten procedures were adequate for bacterial pathogens; bead-beating/ammonium acetate was optimal for microarray detection |
| [76] | Eukaryotes | Freshwater (waste ponds) | UK | commercials kit (3) | Qiagen MoBio | none | Yield Quality | 18S rRNA | NA (PCR) | QIAGEN DNeasy Blood & Tissue yielded the highest DNA yield and quality for freshwater microalgae, while diversity profiles were comparable across kits |
| [77] | Bacteria | Freshwater (River) | China | phenol-chloroform variants (3) | NA | Lysis Purification | Yield quality | 16S rRNA | NA (PCR) | Modified protocol was most efficient, yielding high-quality DNA suitable for molecular applications |
| [78] | Bacteria Eukaryotes | Freshwater (Lake,River) | Switzerland | commercial kits (2)<br>phenol–chloroform (1) | Qiagen MoBio | none | Yield | 16S rRNA COI | metabarcoding targeted | Extraction protocol choice alters biodiversity detection, affecting DNA yield, sequence recovery, and community composition. |
| [24] | Bacteria | Freshwater (River)<br>Wastewater | USA | commercial kits (4) | Qiagen MoBio | none | NA | 16S rRNA | metabarcoding | DNA Mini Kit and PowerSoil Kit yielded the most consistent sequencing results across water and wastewater samples |
| [79] | Prokaryotes<br>Phytoplankton<br>Vertebrates | Seawater (Pacific) | USA | commercial kit (2)<br>phenol-chloroform (1) | Qiagen MoBio | none | Yield | 16S rRNA<br>18S rRNA<br>12S rRNA | metabarcoding | DNeasy and MoBio kits produce comparable results for multi-trophic eDNA, while phenol-chloroform significantly alters community composition |
| [80] | Vertebrates | Freshwater (Stream and aquaria)<br>Tap water | Australia | commercial kit (2) | Qiagen MoBio | none | Yield | 12S rRNA | NA (qPCR) | Extraction with the Qiagen DNeasy kit provided the highest DNA yield and best cost-efficiency, especially when paired with cellulose nitrate filtration. |
| [81] | Bacteria<br>Fungi, Plant<br>Vertebrates,<br>Animal | Freshwater (Stream) | New Zealand | commercial kits (6) | Qiagen MoBio<br>ZymoBIOMICS | none | Yield Quality | 16S rRNA<br>ITS2, 12S rRNA<br>COI | metabarcoding | DNeasy PowerSoil performs best across taxa and sample types, but no single method is optimal for all organisms |
| [82] | Bacteria | Freshwater (Lake) | Brazil | commercial kit (1)<br>phenol–chloroform (1) | MoBio | none | NA | 16S rRNA | metabarcoding | Phenol–chloroform and kit extractions yielded microbial communities too different to compare. |
| [83] | Bacteria Eukaryotes | Freshwater (Reservoir) | China | commercial kits (5) | Qiagen MoBio<br>MP Biomedicals | none | Yield Quality | 16S rRNA<br>18S rRNA | metabarcoding | Extraction methods bias rare taxa more than abundant ones; some protocols favor diversity, others reproducibility |
| [84] | Eukaryotes | Seawater (Atlantic) | Spain | commercial kit (3)<br>Other chemical methods (2) | Qiagen<br>Omega Bio-tek<br>Epicentre | none | Yield Quality | 18S rRNA | metabarcoding | Extraction method choice critically affects detection of difficult-to-identify nano- and picoplankton eukaryotes |
| [85] | Prokaryotes | Seawater (Adriatic Sea) | Croatia | commercials kit (2)<br>phenol-chloroform variants (3)<br>direct PCR (1) | Qiagen | Lysis | Yield | 16S rRNA | metabarcoding | All methods showed similar biases; methodological consistency matters more than method choice |
| This study | Prokaryotes | Freshwater (Lake) | France | commercial kits (18) | Qiagen MoBio<br>Macherey-Nagel<br>Zymo<br>MP Biomedicals | Lysis Elution | Yield | 16S rRNA | metabarcoding | Modified commercial kit increased DNA yield, but higher yield negatively correlated with read count and ASV richness |

**Table 3:** Summary of DNA extraction protocols and performance metrics. Cells are coded as “+” (more favorable) or “−” (less favorable). Handling time was classified as more than 3 hours plus overnight incubation (“--”), more than 3 hours (“-”), approximately 2 hours 30 minutes (“o”), or less than 2 hours 30 minutes (“+”). Cost per extraction was classified as more than €40 (“--”), between €6 and €15 (“o”), or less than €6 (“+”). Contamination was assessed by absorbance ratios A260/230 and A260/280 (“+” indicates at least one ratio close to optimal values: ∼2.0 for A260/230 and ∼1.85 for A260/280; “-” indicates both ratios were not optimal), and fragment length was assessed by gel electrophoresis (“+” indicates high molecular weight DNA ∼50 kb, “-” indicates fragmented DNA 10–15 kb). NA indicates out-of-range data due to low DNA concentration and contamination leading to unreliable ratios. For DNA yield (quantity and reproducibility), ASV richness, and intra-replicate dissimilarity (wUniFrac), categories were defined by tertiles from the first sampling (n = 45). Lower wUniFrac indicates higher compositional reproducibility. For the second sampling (n = 12), categories for E1E, E1F, and E1G were assigned relative to E1D and its tertile from the first sampling.

| Description |  |  | Logistic |  | DNA Yields |  | DNA quality |  | Community Structure |  |
| --- | --- | --- | --- | --- | --- | --- | --- | --- | --- | --- |
| Brand | Kit | Protocol | Handling time | Cost | Quantity | Reproducibility | Contamination | Length | Richness (ASV) | Intra-replicate dissimilarity |
| Experiment K (45 samples) |  |  |  |  |  |  |  |  |  |  |
| Qiagen | DNeasy PowerLyzer PowerSoil | E1A | **o** | **o** | - | + | - | + | + | **o** |
|  |  | E1B | - |  | **o** | + | - | + | **o** | **o** |
|  |  | E1C | - |  | + | + | + | + | - | + |
|  |  | E1D | -- |  | + | + | + | + | - | + |
|  | DNeasy PowerSoil Pro | E2A | **o** | **o** | **o** | **o** | - | + | **o** | - |
|  |  | E2B | -- |  | + | **o** | - | + | - | **o** |
|  |  | E2C | -- |  | + | **o** | + | + | - | + |
|  | DNeasy PowerSoil | E3A | **o** | NA | **o** | - | - | + | **o** | **o** |
|  |  | E3B | -- |  | + | + | + | + | **o** | + |
|  | DNeasy PowerMax Soil | E4A | - | -- | - | - | NA | NA | **o** | - |
|  |  | E4B | -- |  | **o** | **o** | NA | NA | **o** | + |
| Brand | Kit | Protocol | Handling time | Cost | Quantity | Reproductibility | Contamination | Length | Richness (ASV) | Intra-replicate dissimilarity |
| Macherey-Nagel | Nucleospin Soil | E5A1 (Buffer SL1) | 0 | + | - | - | - | NA | + | - |
|  |  | E5A2 (Buffer SL2) | 0 |  | - | 0 | - | NA | + | - |
| Zymo | ZymoBIOMICS DNA | E6A | 0 | 0 | - | - | NA | + | 0 | - |
| MP Biomedicals | FastDNA SPIN | E7A | 0 | 0 | 0 | - | NA | + | + | 0 |
| Experiment L (12 samples) |  |  |  |  |  |  |  |  |  |  |
| Qiagen | DNeasy PowerLyzer PowerSoil | E1D | - - | 0 | + | + | - | + | - | + |
|  |  | E1E | + |  | + | + | + | - | - | + |
|  |  | E1F | + |  | + | + | + | - | - - | + |
|  |  | E1G | + |  | + | + | + | - | - | + |

### DNA yield (quantity and replicability) and community composition

In contrast to large between-kit differences in other matrices such as soil or marine sediments [15,19], all DNA yields were broadly comparable across manufacturer protocols, the NucleoSpin Soil kit showing the lowest, although not significantly different, yields (Fig 1A, Table 3). . The lower yield obtained with the NucleoSpin Soil kit may be explained by several factors, including weaker mechanical or chemical lysis, buffer chemistry that promotes inhibitor carryover or reduces DNA binding, or lower binding/elution efficiency. However, because the exact buffer compositions and lysis chemistries are proprietary and not disclosed, we were unable to distinguish the underlying mechanisms in this study. Interestingly the lower DNA yield of the NucleoSpin Soil kit (E5A1-2, Table 3) was accompanied by higher Evenness, suggesting reduced dominance rather than increased richness (Fig 2A).

Among the Qiagen kits tested, the DNeasy PowerLyzer modified protocol (E1D) yielded up to ∼4× more DNA than the standard protocol (E1A) (Fig 1A, Table 3). These modifications likely acted through complementary mechanisms: freeze-thaw cycles disrupt cells through heat shock [51,52]; incubation at 65 °C enhances chemical lysis and destabilises cell walls and membranes [53–55]; and complete ethanol removal improves DNA desorption from the silica membrane and prevents PCR inhibition. A two-step elution further increases recovery by rehydrating the matrix twice and collecting DNA that remains after the first pass [53,56]. Although the modifications were applied as a bundle, preventing isolation of individual effects, their combined impact enhanced DNA yield. In addition, two other modifications, tested independently, also increased DNA recovery: overnight incubation with agitation in the PowerBead Solution and heating the elution buffer to 65 °C. Overnight agitation likely promoted cell detachment from particulate matter and improved sample homogenization, thereby enhancing the efficiency of bead-beating [57]. Likewise, preheating the elution buffer increased recovery [58], probably by improving membrane rehydration, increasing DNA solubility, and weakening DNA–silica interactions. The improved A260nm/230nm ratios observed with protocol modifications confirm that additional lysis and purification steps effectively reduce contaminants such as carbohydrates and phenolic compounds, which are common in freshwater samples with high particulate matter. However, these modifications also reduced rare ASV detection, reinforcing the trade-off between DNA purity and preservation of low-abundance community members. Improving contaminant removal could degrade DNA from rare ASV, compromising the completeness of community characterization despite higher overall DNA yields. An additional explanation is a better recovery of dominant taxa under stronger lysis. In both experiments, the dominant genera belonged to phyla known for thick cell envelopes, such as Actinobacteriota [59] and Cyanobacteriota [60], which require harsher disruption; their improved recovery would increase their relative abundance and, in turn, lower evenness and richness. To note that, since no mock community was included in this study, all observations remain comparative and do not refer to the “true community composition”.

Within the PowerLyzer-based protocols, all modifications were associated with fewer observed ASVs despite higher DNA yields (Fig 3, Table 3). ASV counts and read depth tended to decrease as total DNA increased, in line with previous observations [61]. Because a fixed volume of extract was used as PCR template rather than a normalised DNA mass, high-yield extracts contributed more template to the reaction; the lower richness they nonetheless yielded therefore cannot be attributed to a smaller amplified fraction, and instead points to extraction-related effects. This negative correlation can be partly explained by co-extraction of non-bacterial DNA. Additionally, DNA shearing, resulting in smaller DNA fragments, represents a critical consequence of intensive mechanical lysis that compromises PCR efficiency and reduces effective amplicon recovery after quality filtering. Intensive lysis benefits are outweighed by the loss of DNA integrity through fragmentation. Multiple cycles (E1E, E1G) were more effective at enriching dominant taxa (e.g. *Cyanobium*) than a single, more intense cycle (E1F), suggesting that repeated mechanical stress, rather than instantaneous lysis intensity alone, drives shifts toward dominance. Conversely, this single intense cycle detected fewer rare ASV, suggesting over-aggressive lysis disproportionately degrades low-abundance DNA and yielded fewer reads and ASVs overall. These patterns reflect a trade-off: stronger lysis raises the total amount of extracted DNA but can degrade sequencing output, requiring an intermediate optimum.

While protocol modifications reduced rare taxa detection, shared ASVs (common to all compared protocols) accounted for > 93% of total reads, suggesting consistent detection of abundant taxa across protocols, with differences concentrated in rare ASVs as shown in previous study [62]. Although the PCoA of the communities of all samples tested did not show any clear pattern in phylogenetic structure, community composition differed significantly among protocols and manufacturers, consistent with previous findings that extraction methods influence the observed prokaryotic community [63,64]. The distinct clustering of PowerMax (E4) samples (Fig 6) likely reflects a mismatch between kit design and sample type: PowerMax is designed to process up to 10 g of soil, whereas other kits handle up to 0.25 g. Water membrane samples represent far lower biomass than these capacities, potentially underloading the PowerMax columns and reducing DNA binding efficiency, which may have altered the representation of certain taxa.

Cell wall and membrane architecture can influence the efficiency of DNA extraction [65,66]. Previous studies highlighted that some protocols preferentially recover Gram-negative taxa [63], whereas others favor Gram-positive taxa [67]. We hypothesised that adding lysis steps would increase the recovery of Gram+ bacteria because their thicker peptidoglycan layers require harsher disruption or other microbes with robust membranes. Contrary to this expectation, several Gram+ genera—*Clostridium*, *Mycobacterium*, *Fictibacillus*, *Tumebacillus*, and *Romboutsia*—were relatively more abundant under the manufacturer protocols than under the modified ones (Fig S5A) [68–72]. A plausible explanation is a relative abundance artefact caused by preferential lysis of Gram-cells. Intensified mechanical and thermal disruption may disproportionately increase the recovery of Gram-DNA, thereby lowering the relative abundance of Gram+ taxa, even if their absolute recovery also increases. In other words, more aggressive lysis may improve Gram+ lysis, but the concurrent gain for Gram-taxa can mask that effect in relative abundance data.

In addition to differences in DNA yield and quality, the time and cost of all 18 protocols were systematically evaluated given their practical relevance (Table 3; Table S2). For some pairs, total processing time varied by roughly 1.8-fold; for example, E1F required about 2h, whereas E1D last for about 4h, excluding the overnight incubation. E4B was the clear outlier, with approximately 6 h of processing plus an overnight incubation and a much higher price, about 300% higher than the other kits. Among the protocols, DNeasy PowerMax Soil (E4) had the highest cost per sample (Table 2-3). This kit has been used for water samples [73] and has been reported as effective for soils [74], but it is designed for large input masses. This requires larger buffer volumes and higher-capacity columns, which increases cost. In addition, E4 is sold only in boxes of 10 extractions, whereas other kits are available in larger formats (up to 250), further increasing the per-sample cost. Among the standard and modified versions of DNeasy PowerMax Soil, the modified protocol (E4B) had a longer processing time, mainly because the freeze-thaw phase required more time at each step.

Concluding, the fundamental challenge that emerged here is the balance between recovering sufficient and good quality DNA versus effects on downstream sequencing output (reads and ASVs) (Table 3). Our results suggested that commercial DNA extraction kits, and protocol modifications - of lysis and elution - can affect both yield and the inferred prokaryotic community composition in freshwater samples. These effects were kit-specific: for example, intensifying the PowerLyzer workflow increased total DNA, whereas varying FastPrep settings did not change yield but did alter diversity patterns. Because we evaluated a single freshwater matrix, the magnitude and direction of these effects may differ in other matrices [64]. Choosing a protocol is therefore a multi-criteria decision that balances logistics (equipment, hands-on time, cost) against a clear trade-off between total DNA recovered and downstream sequencing output (reads and ASVs) (Table 3). In our data, read counts and ASV richness tended to decrease as total DNA increased. If high yield and reproducibility are the primary goals and longer handling is acceptable, PowerLyzer with all modifications (E1D) is a good compromise, while stronger mechanical lysis (e.g E1E) provides similar yield with less hands-on time. However, both settings reduced ASV richness and read counts, so if the priority is maximizing usable reads, unmodified manufacturer protocols are preferable, which also preserve DNA integrity for long-read sequencing.

Finally, these findings can be complemented by additional tests, to better distinguish DNA recovery, integrity, and downstream performance: (i) explicitly test for PCR inhibition using dilution series and internal spike-ins, and consider post-extraction cleanup; (ii) include absolute quantification (e.g. qPCR or ddPCR) targeting total 16S rRNA genes and selected taxa to evaluate how lysis conditions affect the detection of low-abundance groups.

Last but not least, these findings are of primary importance for metagenomic-based studies of microbial communities since for this approach, high DNA quantities are usually required to process the sequencing step.

## Acknowledgements

We thank P. Bertolino for laboratory assistance with DNA extraction and amplification procedures. We acknowledge PRISMARCTYC project (ANR-21-SOIL-0003), the graduate school IFSEA (ANR-21-EXES-0011) and the CPER-IDEAL (2022-2027). A. Szylit was funded by a Ph.D. grant from the ‘Région Hauts-de-France’ and the ‘Université du Littoral Côte d’Opale (ULCO)’.

## Data availability statement

Sequence data have been deposited in the European Nucleotide Archive under project accession number PRJEB94317.

The complete R pipeline (v1.0.0) and datasets supporting this study are available on Zenodo: https://doi.org/10.5281/zenodo.18421569

## Author contributions

Conceptualization: A.S., L.J., MC.C. Methodology: A.S., MC.C. Investigation: A.S. Data curation: A.S. Writing – original draft: A.S. Writing – review and editing: A.S., L.J., MC.C., L.C., M.B., U.C. Supervision: L.J., M.B., L.C., U.C. Funding acquisition: U.C., L.J. All authors have read and agreed to the published version of the manuscript.

