## Supplementary material for "Comparing DNA Extraction Protocols for Freshwater Prokaryotic Communities: Impacts on Yield and Microbial Profiling"

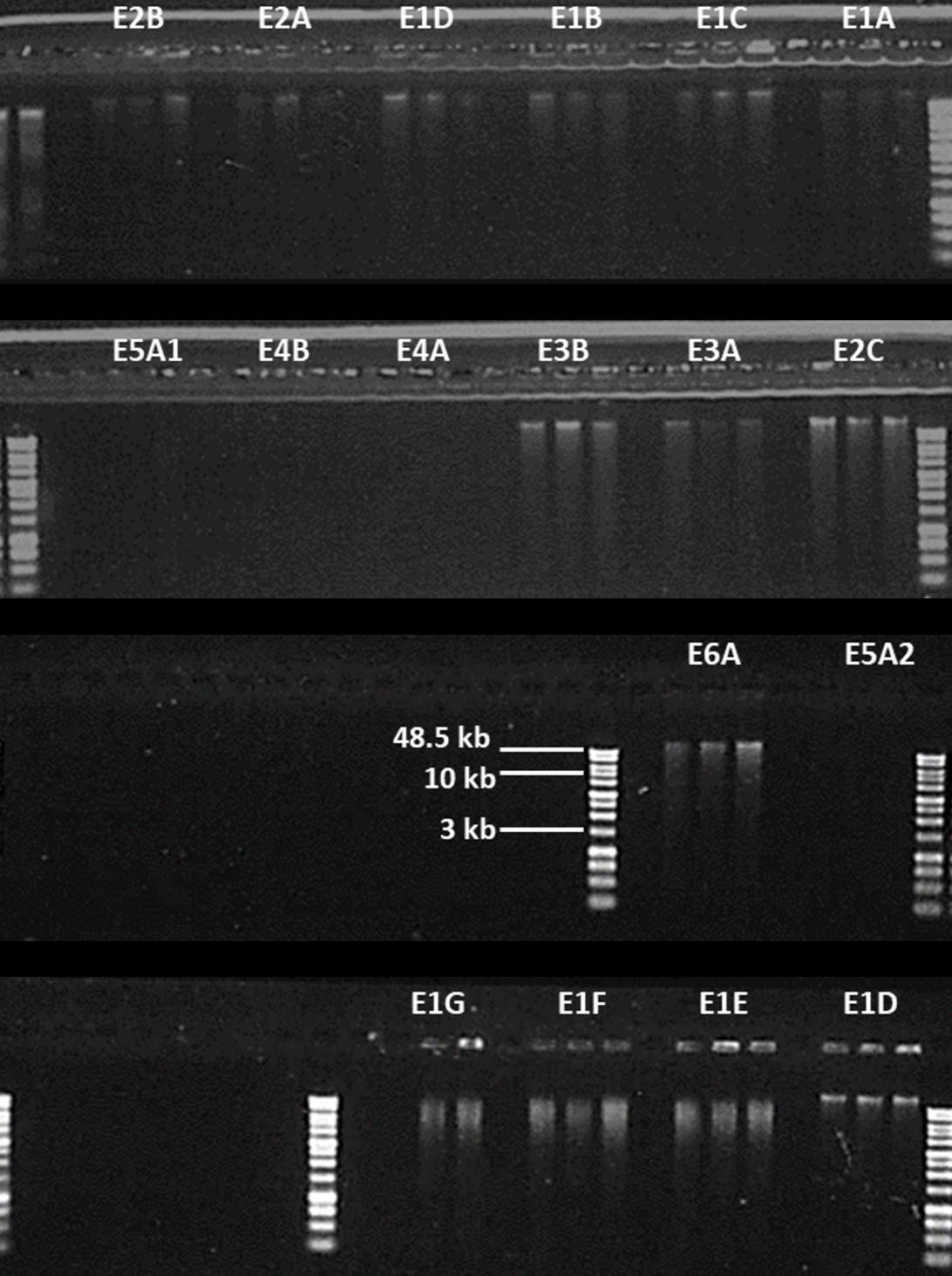

Figure S1: Agarose gel electrophoresis showing DNA fragment size for extraction protocols in both experiments. Each triplicate was loaded on the gel. No detectable signal was observed for E4 and E5. The molecular weight marker is the Quick-Load® 1 kb Extend DNA Ladder (New England Biolabs).

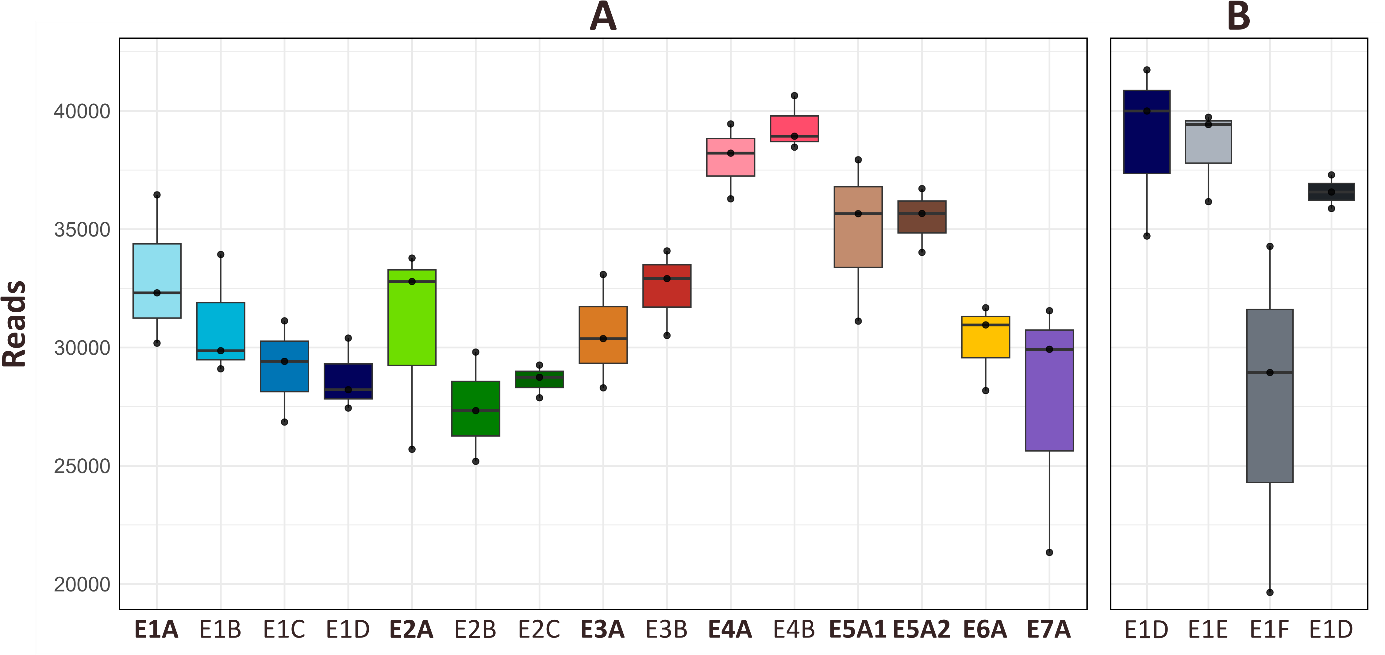

Figure S2: Sequencing reads obtained after filtering (removal of mitochondria, chloroplasts, and singletons) across extraction protocols during the first (A) and second (B) sampling. Protocols in bold correspond to manufacturers' protocols. Panels A and B now share the same y-axis scale.

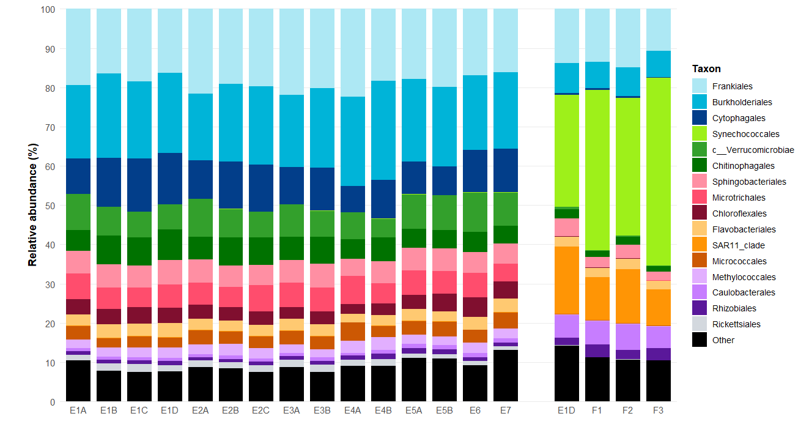

Figure S3: Composition of the 16 most abundant orders across extraction protocols. Taxonomic assignment is based on the lowest assigned rank, with prefixes (e.g. C_) indicating classification at higher taxonomic levels (Class) when order-level assignment was unavailable. Percentages on the x-axis represent relative abundance within the total prokaryotic community. 'Other' includes Order representing < 1.1% of total reads. Protocols in bold correspond to manufacturers' protocols

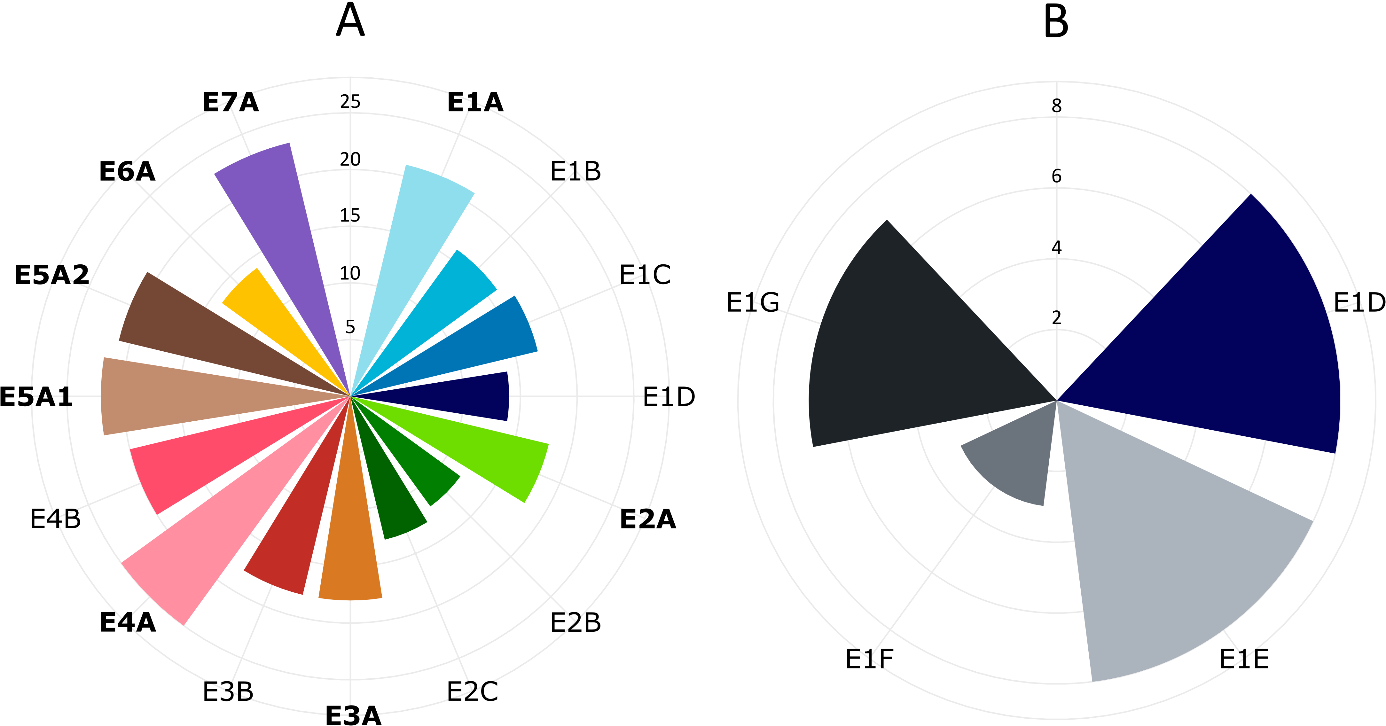

Figure S4: Cumulative abundance of rare taxa (<0.01%) for Experiment K (A) and Experiment L (B). Values are summed across triplicates.

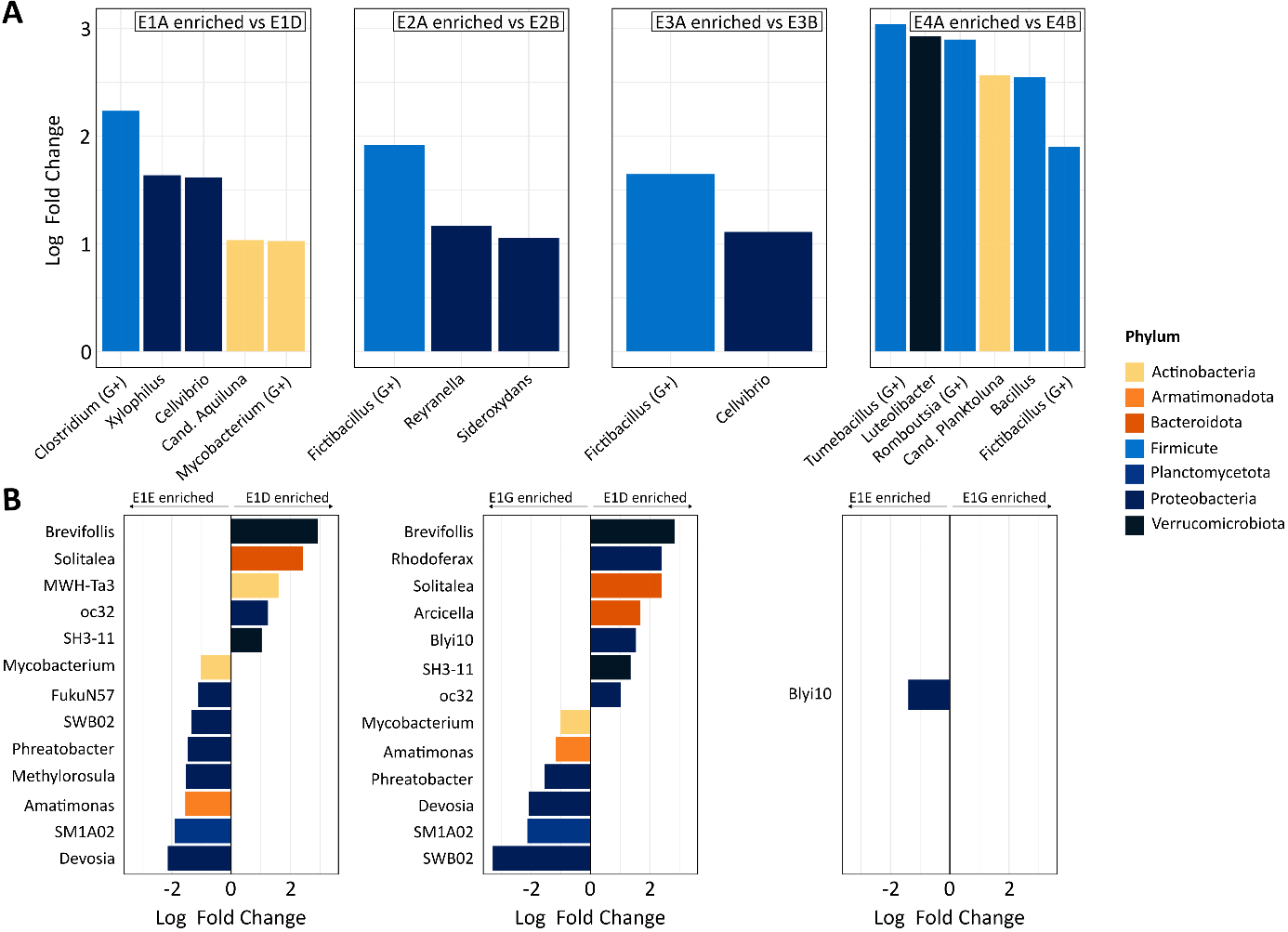

Figure S5: Differential abundance analysis of genera using ANCOM-BC (log2 fold change threshold = 1). (A) Genera significantly enriched in standard protocols compared to modified protocols for each Qiagen kit during for Experiment K (E1 to E4). (B) Genera significantly enriched between protocols for Experiment L. Comparisons involving E1F are provided in Table S3. Only genera meeting the significance threshold are shown. Colors indicate the corresponding phylum. G+ denotes Gram-positive genera.

Table S1: Components Details for Polymerase Chain Reaction (PCR) and thermocycling procedure The components used for the Polymerase Chain Reaction (PCR), including the (A) master mix and the (B) PCR mix, were carefully prepared to ensure sterility and precision. Nuclease-free water was UV-treated for 45 minutes before each use. The 10X buffer, MgCl₂, and Taq polymerase came from the Platinum Taq DNA Polymerase Kit (Invitrogen). The dNTPs were obtained from the PCR Nucleotide Mix (Promega), while the DNase and DTT came from the PCR Decontamination Kit (enzo). A fixed volume of 1 µl of DNA extract was added to each reaction, regardless of its concentration. All reagents were used at the concentrations recommended by the Platinum Taq DNA Polymerase Kit (Invitrogen) and the Earth Microbiome Project 16S protocol.

| **Components** | **Volume (µL)** |
| --- | --- |
| **Master Mix** | **22.75** |
| **Primer Fwd** | **1** |
| **Primer Rev** | **1** |
| **Taq** | **0.2** |
| **DNase** | **0.63** |
| **DTT** | **0.63** |

**A	B**

| **Components** | **Volume (µL)** |
| --- | --- |
| **H_2_O** | **19** |
| **Buffer 10X** | **2.5** |
| **MgCl_2_** | **0.75** |
| **dNTP** | **0.5** |

Table S2 : Summary of processing time and cost for DNA extraction protocols. Prices were retrieved from manufacturers’ websites on 25 July 2025 and correspond to list prices at that date, excluding taxes and shipping.

| Brand | Kit | Protocol | Approximate processing time (12 samples) | Price per extraction (€) |
| --- | --- | --- | --- | --- |
| **Experiment K ( 45 samples)** | | | | |
| Qiagen | DNeasy PowerLyzer PowerSoil | E1A | 2h30 | 8.90 |
|  |  | E1B | 3h50 |  |
|  |  | E1C | 3h50 |  |
|  |  | E1D | 3h50 (+ 1 night) |  |
|  | DNeasy PowerSoil Pro | E2A | 2h30 | 10.20 |
|  |  | E2B | 3h50 (+ 1 night) |  |
|  |  | E2C | 3h50 (+ 1 night) |  |
|  | DNeasy PowerSoil | E3A | 2h30 | Not available discontinued |
|  |  | E3B | 3h50 (+ 1 night) |  |
|  | DNeasy PowerMax Soil | E4A | 3h15 | 40.20 |
|  |  | E4B | 6h (+ 1 night) |  |
| Macherey-Nagel | Nucleospin Soil | E5A1 (Buffer SL1) | 2h30 | 5.72 |
|  |  | E5A2 (Buffer SL2) | 2h30 |  |
| Zymo | ZymoBIOMICS DNA | E6A | 2h30 | 8.24 |
| MP Biomedicals | FastDNA SPIN | E7A | 2h30 | 13.84 |
| **Experiment L (12 samples)** | | | | |
| Qiagen | DNeasy PowerLyzer PowerSoil | E1D | 3h50 | 8.90 |
|  |  | E1E | 2h10 |  |
|  |  | E1F | 2h05 |  |
|  |  | E1G | 2h10 |  |

Table S3: Differential abundance analysis of genera using ANCOM-BC between E1F and the 3 other protocols from the second sampling date

| **Comparison** | **Genus** | **Enriched in** | **Log fold change** |
| --- | --- | --- | --- |
| E1D vs F2 | Brevifollis | E1D | 2.5 |
|  | Solitalea |  | 2.2 |
|  | Arcicella |  | 1.4 |
|  | MWH-Ta3 |  | 1.4 |
|  | Oc32 |  | 1.2 |
|  | SH3-11 |  | 1.1 |
| F1 vs F2 | Flavobacterium | F1 | 1.3 |
|  | Armatimonas |  | 1.2 |
| F2 vs F3 | Hydrogenophaga | F2 | 2.6 |
|  | SWB02 | F3 | 2.9 |
